# Local administration of bicuculline into the ventrolateral and medial preoptic nuclei modifies sleep and maternal behavior in lactating rats

**DOI:** 10.1101/2021.04.08.439015

**Authors:** Luciana Benedetto, Mayda Rivas, Florencia Peña, Diego Serantes, Annabel Ferreira, Pablo Torterolo

## Abstract

The preoptic area (POA) is a brain structure classically involved in a wide variety of animal behavior including sleep and maternal care. In the current study, we evaluate the specific effect of disinhibition of two specific regions of the POA, the medial POA nucleus (mPOA) and the ventrolateral POA area (VLPO) on sleep and maternal behavior in lactating rats. For this purpose, mother rats on postpartum day 1 (PPD1) were implanted for polysomnographic recordings and with bilateral cannulae either in the mPOA or in the VLPO. The rats were tested for sleep and maternal behavior on PPD4-8 after the infusion of the GABA-A antagonist, bicuculline (0, 10 or 30 ng/0.2 µl/side). Infusion of bicuculline into the mPOA augmented retrieving and nest building behaviors and reduced both nursing and milk ejections but had almost no effect on sleep. When bicuculine was microinjected into the VLPO, the rats significantly increase the number of retrievings and mouthings and reduced the nursing time without changes in milk ejections, which was associated with an increase in wakefulness and a reduction in light sleep.

Our results show that disinhibition of the mPOA, a key area in the control of maternal behavior, increased active maternal behaviors and reduced nursing without affecting wakefulness or sleep time. In contrast, the enhancement of some active maternal behaviors when the drug was infused into the VLPO, a sleep-promoting area, with a concomitant increase in wakefulness suggests that mother rats devote this additional waking time in the active maternal care of the pups. We hypothesize that maternal behavior changes after bicuculine microinjection into the VLPO is caused by a reduction in the sleep drive, rather than a direct effect on maternal behavior.

## Introduction

The preoptic area (POA) has been recognized as the key sleep-promoting center almost one century ago [1]. Within this area, both the medial preoptic area (mPOA) [2, 3] and especially the ventrolateral preoptic area (VLPO) [4, 5] have been extensively related to the generation and maintenance of sleep. However, the role of these areas in the control of sleep seems to differ [6-8]. For instance, Sryvidya et al. (2006) reported that while the destruction of neurons within the mPOA reduces both NREM and REM sleep, the same lesions in VLPO only affect NREM sleep in male rats [6]. Since the brain of the mother rat, and specifically the POA, undergoes great anatomical and functional changes throughout the postpartum period [9-13], it could be speculated that the role of the mPOA and the VLPO on sleep and waking behavior may change during this period.

The mPOA has been postulated as the key component in the coordination and integration of maternal behavior in virtually all mammals [14]. The mPOA contains receptors for hormones, neurotransmitters and neuromodulators involved in the rapid onset of maternal behavior after parturition [15-19]. While other brain areas have also been implicated in the control of maternal behavior, as far as we know, there are no studies that evaluated the role VLPO in this behavior.

It has been reported a close relationship between maternal behavior and sleep [20]. Lactating rats exhibit a unique pattern of sleep [21], due in part to accomplish the maternal care of the pups. In this sense, we showed that rats mostly sleep while nursing [22]. Thus, it would be possible to speculate that sleeping areas could indirectly, through promoting or reducing sleep, modify maternal behavior.

The POA contains neurons that use GABA as a neurotransmitter and expresses GABA-A receptors that has been related to sleep [7, 8, 23-25]. In this sense, Mendelson et al. have shown that triazolam, a GABA-A agonist, promotes NREM sleep within the mPOA [8], but this drug at the same dose did no alter sleep when microinjected into the VLPO of male rats [7]. On the other hand, Arrati et al (2006) show that the GABA-A receptor agonist, muscimol, injected in the mPOA, produced dose-dependent deficits in most active components of maternal behavior [26].

As microinjections of GABA-A receptor agonists into the mPOA promote NREM sleep in male rats [7, 8], and decrease active maternal behaviors in postpartum rats [26], we hypothesize that bicuculline (BIC), a GABA-A receptor antagonist, may decrease sleep and nursing and promote active maternal behaviors when injected in this area. Despite the lack of studies concerning the role of GABA in VLPO in the control of maternal behavior, given the importance of this nucleus in sleep regulation [4, 5] and the close relationship between sleep and maternal behavior [22, 27], we can speculate that the disinhibition produced by BIC in this area may also decrease sleep and reduce nursing behavior.

In order to explore these hypotheses, in the present report we aimed to determine the effect produced by perfusion of BIC into the mPOA and VLPO on sleep and maternal behavior in lactating rats.

## 2. Methods

### 2.1. Animals and housing

Twenty primiparous Wistar female rats (250-320 g) and their pups were used in this study. All animal use and experimental procedures were in strict accordance with the “Guide to the care and use of laboratory animals” (8th edition, National Academy Press, Washington D.C., 2011) and approved by the Institutional Animal Care Committee. All efforts were made to minimize the number of animals used and their suffering.

Two days before giving birth, pregnant females were housed individually in transparent cages (40 x 30 x 20 cm) containing shredded paper towels as nest-building material. The animals were placed in a temperature-controlled (22 ± 1 °C), sound-proof and electromagnetically shielded recording chamber fitted with slip rings and cable connectors for bioelectrical recordings, under a 12-h light/dark cycle (lights on at 6:00 am), with *ad libitum* access to food and water. On postpartum day 1 (PPD1, birth = day 0), litters were culled to four female and four male pups per mother.

### 2.2. Stereotaxic surgery

The surgical procedures were similar as previous studies of our group [27, 28]. On the morning of PPD1, females were anesthetized with a mixture of ketamine/ xylazine/ acepromazine maleate (80/2.8/2.0 mg/kg, i.p.). Using a stereotaxic device, female rats were bilaterally implanted with a 22-gauge stainless steel guide cannulae with a dummy cannula (Plastic One, Roanoke, VA) aimed 2 mm dorsal to the mPOA: AP −0.5 mm (from Bregma); ML ± 0.5 mm; DV −6.5 mm (from skull surface) or the VLPO: AP −0.5 mm (from Bregma); ML ± 1.0 mm; DV −6.8 mm (from skull surface, according to [29]). In addition, cortical electroencephalogram (EEG) electrodes and dorsal neck muscle electromyogram (EMG) electrodes were implanted for the assessment of sleep and wakefulness (W) states. Recording electrodes for EEG were placed in the frontal cortex (AP = + 4.0; ML = 2.0), parietal cortex (AP = −4.0, ML = 2.0), occipital cortex (AP = −7.0, ML = 3.0), and over the cerebellum as a reference electrode (AP = −11.0, ML = 0.0) [29]. Two additional stainless-steel screws were implanted into the skull as anchors. All electrodes were soldered to a six-pin connector. The connector and the guide cannulae were cemented to the skull using dental acrylic.

A pre-surgery single dose of ketoprofen (5mg/kg, s.c.) was used to reduce pain. In addition, topic antibiotic (Crema 6A, Labyes) was applied into the surgery injury and sterile 0.9 % saline (10 ml/kg, s.c.) was administrated post-surgery to prevent dehydration during recovery.

During surgery, pups were maintained in their home cage under a heat lamp. Immediately after surgery, each mother was reunited with her pups in the home cage located in the recording chamber until the end of the experiments.

### 2.3. Experimental design

Mother rats were randomly assigned to one of two independent groups: VLPO or mPOA treatment. All experiments were performed between PPD4-8 during the light phase. Before experiments began, a baseline recording session was performed to corroborate that all sleep parameters and maternal behaviors were adequate. Each animal received 3 microinjections: 10, 30 ng of BIC and vehicle in a counterbalanced design. The following day after the microinjections, no experiments were performed.

### 2.4. Drug

BIC (Phoenix Pharmaceuticals Inc., Belmont, CA, #070-47) was dissolved in sterile 0.9 % saline to obtain a final concentration of 50 and 150 ng/µl. Aliquots for these doses were prepared in advance, frozen at - 20 °C, and thawed immediately before use.

### 2.5. Microinjection procedure

Females were bilaterally microinjected into the mPOA or the VLPO with either 10 ng of BIC (BIC_10_), 30 ng of BIC (BIC_30_) dissolved in 200 nl of saline, or the same volume of saline over a period of 2 minutes, with the injection cannulae (28 gauge) extending 2 mm beyond the tip of guide cannulae, using a constant-rate infusion pump (Harvard apparatus, USA). The administration cannulae were left in place for an additional minute to allow for the diffusion of the drug. Similar doses were employed in previous studies [30, 31].

### 2.6. Experimental sessions

We used the same experimental protocol in previous studies (for details see [27]). Briefly, pups were removed from the maternal cage for three hours. Before the end of maternal separation, the entire litter was weighed and concomitantly, the microinjection procedure was performed. Immediately after, the mother rat was connected to the recoding system. Once the maternal separation was completed, the pups were scattered in the mothers’ home cage opposite to the nest and the polysomnographic recording and collecting maternal behavior data using a video camera were initiated. After a 4-hour-recodring session, the mother rat was disconnected from the recording device and the entire litter was weighed again [27].

### 2.7. Maternal behavior

Maternal behavior was tested for 30 minutes, as previously described [32]. Accordingly, the number of the following active maternal behaviors measured were: retrieval of the pups to the nest, mouthing (oral re-arrangement of a pup within the nest), licking (corporal and anogenital) and nest building. Also, the latencies to group the entire litter into the nest were measured. In addition, the latencies (first episode ≥ 2 min in duration) and durations of passive behaviors, including hovering over the pups, nursing (kyphotic and supine postures), as well as the total time in contact with the pups (hover over plus nursing) were assessed. In addition, the number of milk ejections was indirectly quantified through the stretching behavior of the pups [33, 34]. The litter weight gain (LWG) was used as an indirect measurement of the amount of ejected milk [27, 35], calculated as the percentage of the difference between the final and initial weight of the entire litter [36].

Any behavior that was not initiated or completed within the 30-min test period was given a latency of 1800 s. The number of eating and self-grooming behaviors was also annotated [27, 32].

### 2.8. Sleep recording

Bioelectric signals were amplified (x1000), filtered (0.1–500 Hz), sampled (1024 Hz, 16 bits) and stored in a PC for further analysis using the Spike 2 software. The states of light sleep (LS), slow wave sleep (SWS), REM sleep and W were determined in 5-seconds epochs with standard criteria [22, 27, 28, 36]. Additionally, the intermediate stage was also distinguished (IS, sleep spindles combined with theta activity [27, 37]).

Total time spent in W, LS, SWS, NREM sleep (LS+SWS), IS and REM sleep over the total recording time and each hour were analyzed. To have a representative analysis of sleep during the maternal behavior test, we also analyzed sleep and W values during the first half hour. In addition, sleep latencies (first episodes ≥ 20 seconds from the beginning of the recordings), number, and duration of episodes of each state were calculated.

Although classical criteria for NREM analysis include LS, SWS and IS, previous data [27] and recent unpublished data from our laboratory strongly suggest that IS should be considered an independent state, and must not be classified as NREM sleep.

### 2.9. EEG spectral power analysis

Digitized, raw EEG signals in prefrontal and parietal channels were exported from Spike2 software into MATLAB at 1024 Hz sampling frequency. The power spectrum was obtained by means of the pwelch built-in function in MATLAB (parameters: window = 1024, noverlap = [], fs = 1024, nfft = 1024), which corresponds to 1-s sliding windows with half-window overlap, and a frequency resolution of 1 Hz [38]. All spectra were normalized to obtain the relative power by dividing the power value of each frequency by the sum across frequencies (total power). Because notch filters were applied to EEG channels in some recordings, power analysis was performed up to 45 Hz. EEG spectral analysis was then conducted during the first half hour, and for the total recording time for each frequency band (δ [1-4 Hz], θ [5-9 Hz], σ [10-15 Hz] β [15-30] and γ [30-45 Hz]. For each animal in each treatment group, the mean power spectrum in each behavioral state (W, LS, SWS, IS and REM) was obtained by averaging the power spectra across all available windows in prefrontal and parietal cortices.

As most animals did not present neither IS nor REM sleep during the first half hour, these states were removed from the analysis in this time-window.

### 2.10. Histological verification of microinjection sites

At the end of the experiment, each animal was euthanized with an overdose of ketamine/ xylazine mixture (i.p.), perfused with 4% paraformaldehyde, and their brains were removed for histological processing. Thereafter, the brains were cut in 150 µm coronal sections with a vibratome. The location of mPOA and VLPO microinjection sites were verified according to [29].

### 2.11. Statistics

Data from maternal parameters did not follow a normal distribution (Kolmogorov-Smirnov test, p < 0.05). Thus, maternal parameters are presented as median ± SIQR (semi-interquartile range) and statistical differences between experimental and control groups were evaluated using Friedman test followed by a post hoc comparison using Wilcoxon signed-rank test. Comparisons between VLPO and mPOA control groups were analyzed using Mann Whitney U test [39].

As sleep parameters and power spectrum of EEG data follow a normal distribution (Kolmogorov-Smirnov test, p > 0.05), values are presented as mean ± S.E.M. (standard error) and the statistical significance of the differences between controls versus drug effects were evaluated utilizing one-way repeated measures (ANOVA) followed by Tukey *post hoc* test. Comparisons between VLPO and mPOA control groups were analyzed using Student T test for independent samples.

The criterion used to discard the null hypotheses in all cases was p < 0.05.

## 3. Results

### 3.1. Sites of injection

As shown in Figure 1A, the microinjection sites in seven rats were located within the mPOA between −0.24 and −0.48 mm from Bregma, based on examination of cannulae tracks in histological sections [29]. Two animals were excluded from the data analysis due to misplaced cannulae. Figure 1B shows the microinjection site of eight rats located within the VLPO. Three rats were excluded from the data analysis due to the cannulae were out of the VLPO borders.

**Figure 1.**
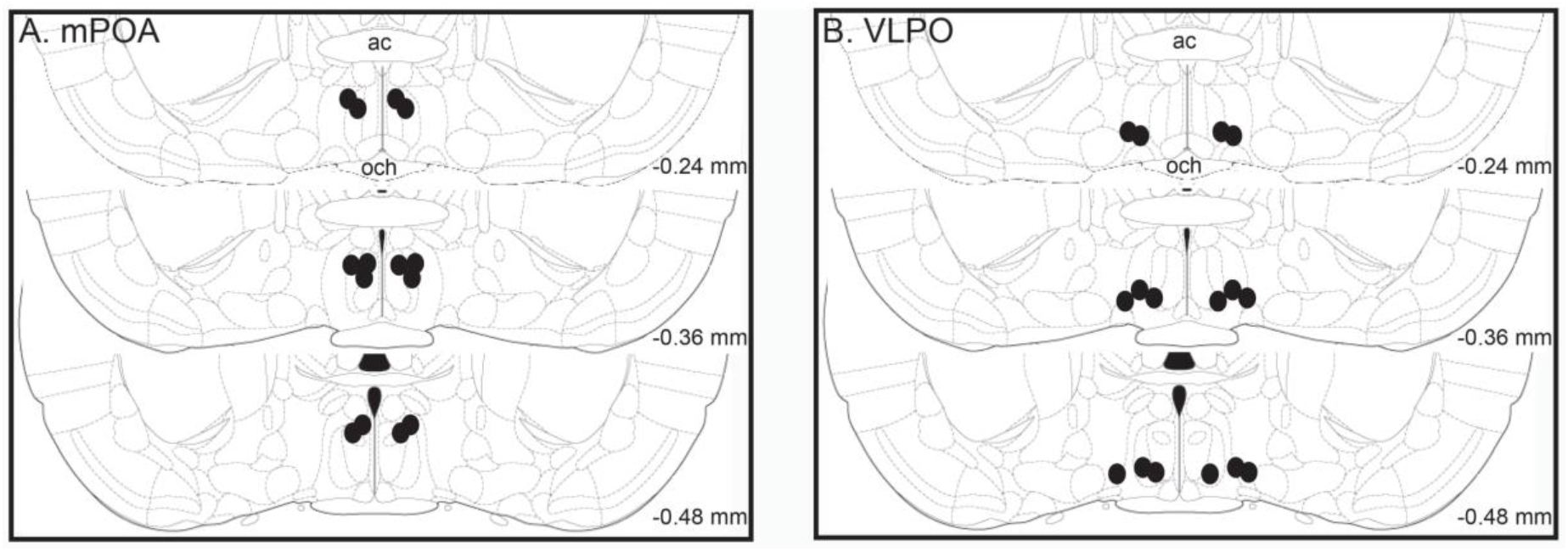
Microinjections sites of bicuculine at the mPOA (A) and VLPO (B). Schematic representations of the microinjection sites in the mPOA (n = 7) and VLPO (n=8) are indicated by black circles. The perfusion sites were recognized by the analysis of the lesions produced by the cannulae. Numbers indicate distance from Bregma. Plates were taken from the atlas of [29]. ac, anterior commissure; och, optic chiasm.

### 3.2. Bicuculline effects on mPOA

#### 3.2.1. Maternal behavior

Maternal behavior components after the microinjection of vehicle, BIC_10_ and BIC_30_ within the mPOA are shown in Figure 2 and Table 1.

**Table 1.**
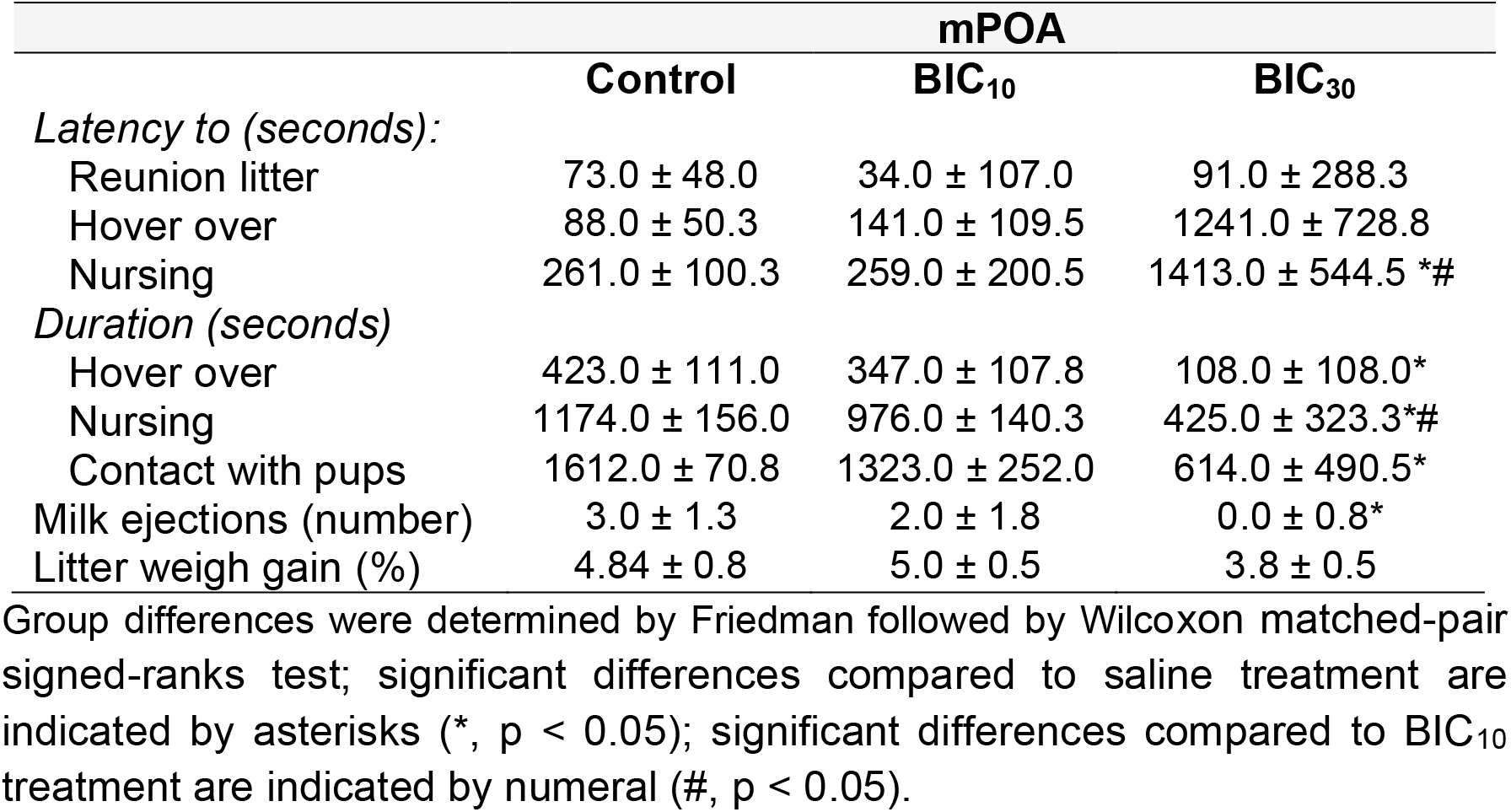
Maternal behavior after BIC and saline microinjections into the mPOA. Effects of bilateral microinjections of saline and BIC (10 and 30 ng/0.2 μl/side) into the medial preoptic area (mPOA) on maternal behavior.

|  | **mPOA** |  |  |
| --- | --- | --- | --- |
|  | **Control** | **BIC<sub>10</sub>** | **BIC<sub>30</sub>** |
| *Latency to (seconds):* |  |  |  |
| Reunion litter | 73.0 ± 48.0 | 34.0 ± 107.0 | 91.0 ± 288.3 |
| Hover over | 88.0 ± 50.3 | 141.0 ± 109.5 | 1241.0 ± 728.8 |
| Nursing | 261.0 ± 100.3 | 259.0 ± 200.5 | 1413.0 ± 544.5 *# |
| *Duration (seconds)* |  |  |  |
| Hover over | 423.0 ± 111.0 | 347.0 ± 107.8 | 108.0 ± 108.0* |
| Nursing | 1174.0 ± 156.0 | 976.0 ± 140.3 | 425.0 ± 323.3*# |
| Contact with pups | 1612.0 ± 70.8 | 1323.0 ± 252.0 | 614.0 ± 490.5* |
| Milk ejections (number) | 3.0 ± 1.3 | 2.0 ± 1.8 | 0.0 ± 0.8* |
| Litter weigh gain (%) | 4.84 ± 0.8 | 5.0 ± 0.5 | 3.8 ± 0.5 |

Group differences were determined by Friedman followed by Wilcoxon matched-pair signed-ranks test; significant differences compared to saline treatment are indicated by asterisks (\*, $p < 0.05$ ); significant differences compared to BIC<sub>10</sub> treatment are indicated by numeral (#, $p < 0.05$ ).

**Figure 2.**
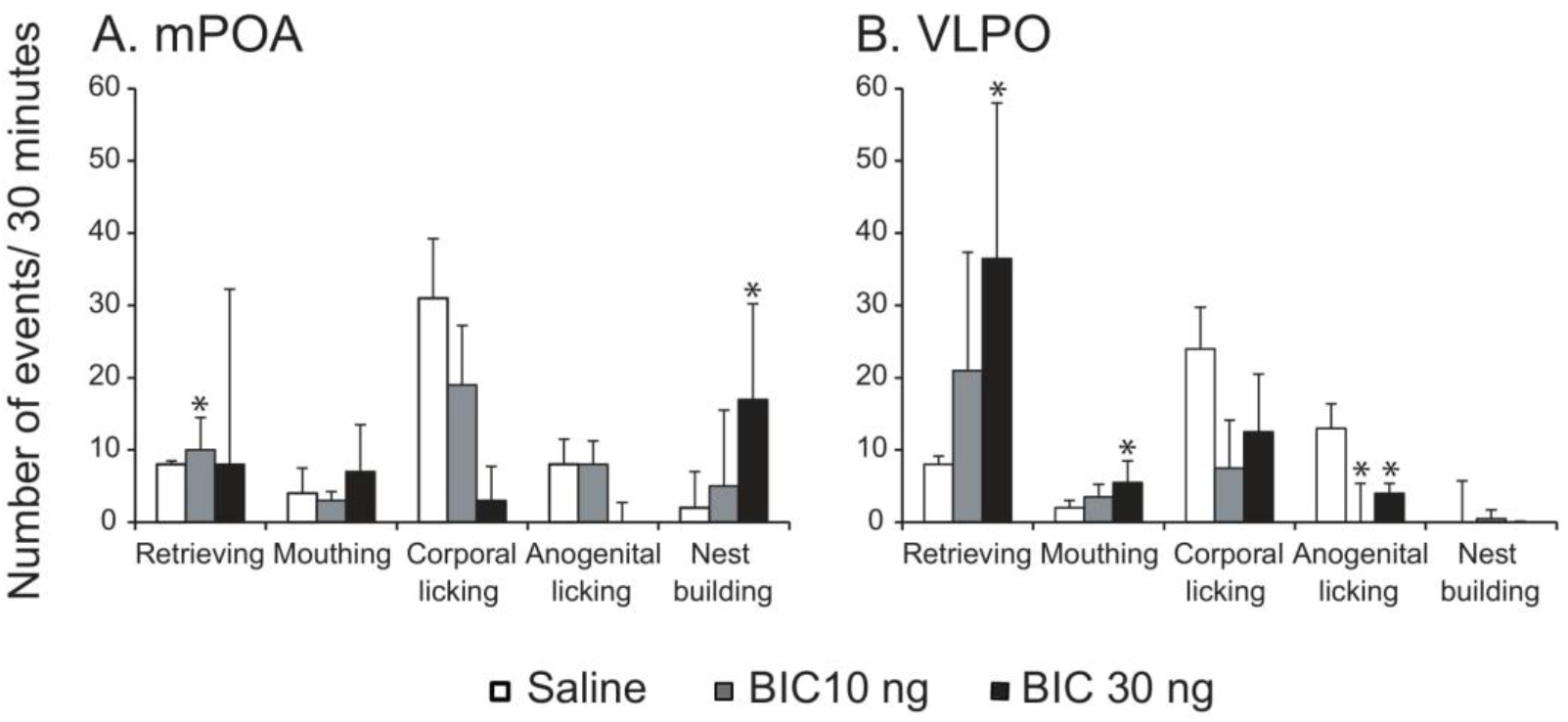
Effects of bicuculline (BIC) microinjections into the mPOA (A) and VLPO (B) on active components of maternal behavior. Graphic charts show the median ± SIQR (semi-interquartile range) of the number of different active maternal responses in a 30-minute-maternal test after bilateral administration of saline and BIC (10 and 30 ng) into the mPOA and VLPO. Intra-group differences were determined by Friedman followed by Wilcoxon matched-pair signed-ranks test; significant differences compared to control group are indicated by asterisks (*, p < 0.05).

The microinjection of BIC_10_ into the mPOA significantly increased the number of pups retrieved to the nest in comparison with the infusion of saline (T_7_ =1.0, p = 0.046), whereas BIC_30_ increased the number of nest building (T_7_ = 1.0, p = 0.027, see Figure 2). None of the other active maternal behaviors were affected by the drug (see Figure 2).

Compared to saline microinjections, BIC_30_ decreased the time total time in contact with the pups (T_7_ = 0.0; p= 0.017), reduced the time hovering over the pups (T_7_ = 1.0, p = 0.027) and the number of milk ejections (T_7_ = 0.0; p = 0.027). In addition, BIC_30_ decreased the time the females nurse their pups compared to control group (T_7_ = 0.0; p = 0.017) and to the lowest dose (T_7_ = 2.0; 0.042, see Table 1).

The highest dose of BIC increased the latency to begin nursing compared to saline microinjection (T_7_ = 1.0, p = 0.027) and to the lowest dose (T_7_ = 1.0, p = 0.027, see Table 1).

Non-maternal behaviors during the maternal behavioral are shown in Supplementary Table S1. Neither eating nor self-grooming were significant different among groups.

#### 3.2.2. Sleep

Statistical comparisons were non-significant for most sleep parameters (see Figure 3 and Table 2), except for the total IS recording time that was significantly different among groups (F (2, 12) = 3.947, p = 0.048); post hoc analysis did not reach significant differences, but time in IS tended to be higher after BIC_10_ treatment compared to saline (p = 0.082) and BIC_30_ (p = 0.069). Additionally, when analyzed the first half hour and each hour independently, only the time spent in REM sleep during the second hour differ among groups (F (2, 12) = 4.255, p =0.040), specifically reduced in BIC_30_ compared to control treatment (p = 0.032). The number of episodes, frequency and its duration, as well as latencies for each behavioral stage remained unchanged among groups (see Table 2).

**Table 2.**
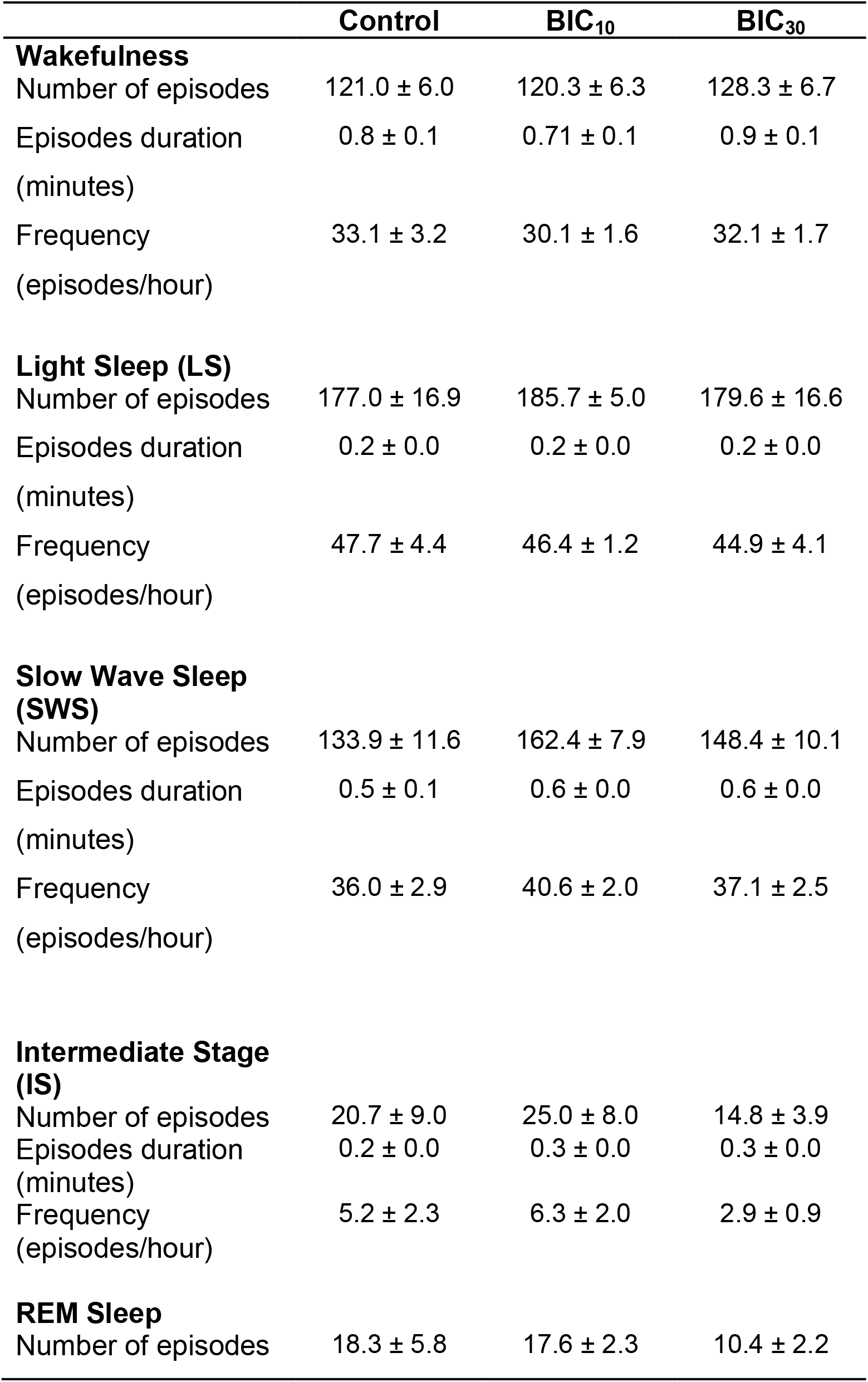
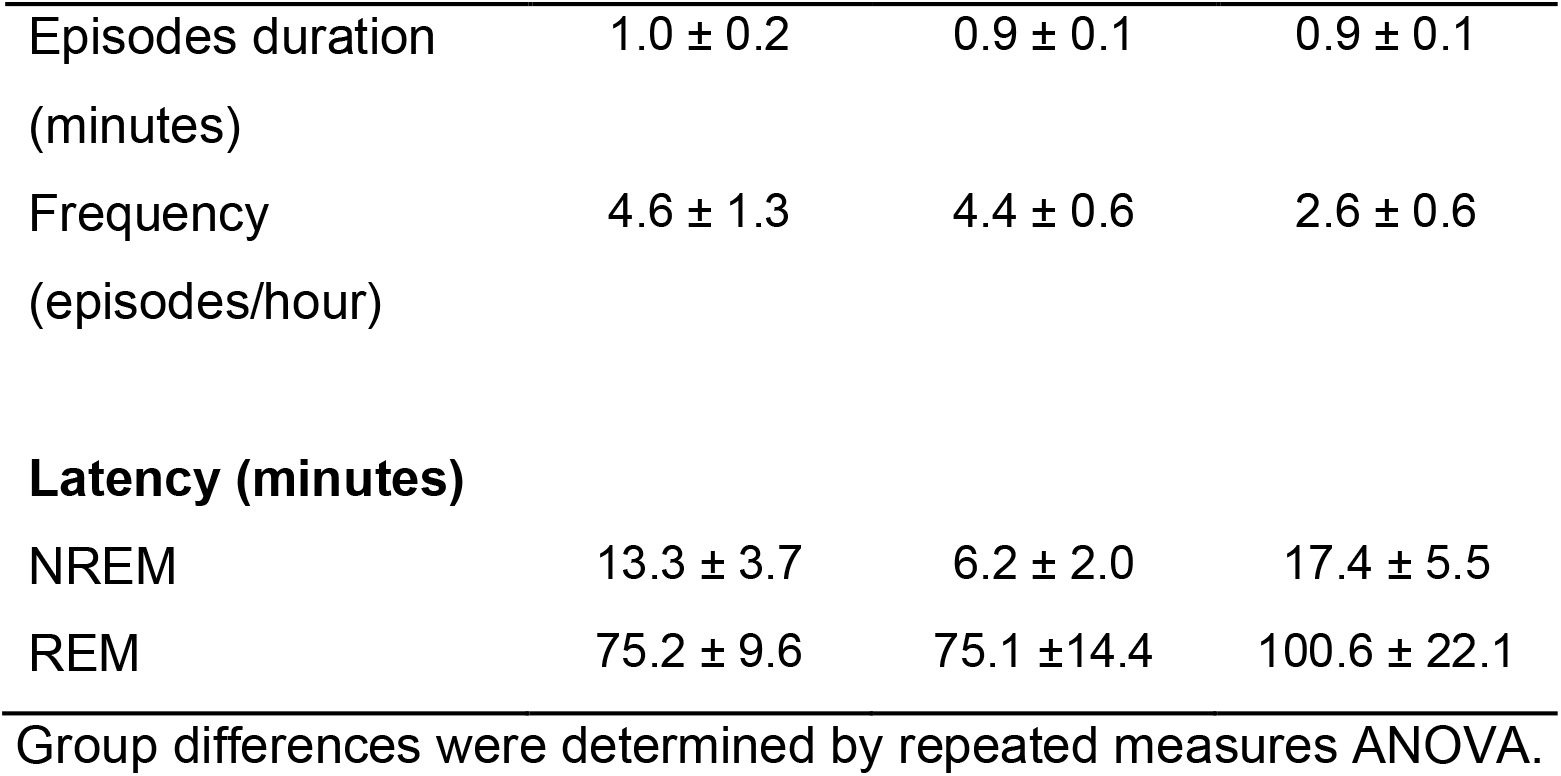
Sleep and waking effects after bicuculline (BIC) and saline microinjections into the medial preoptic area (mPOA). Effects of bilateral microinjections of saline and BIC (10 and 30 ng/0.2 μl/side) into the mPOA on sleep parameters.

|  | **Control** | **BIC<sub>10</sub>** | **BIC<sub>30</sub>** |
| --- | --- | --- | --- |
| **Wakefulness** |  |  |  |
| Number of episodes | 121.0 ± 6.0 | 120.3 ± 6.3 | 128.3 ± 6.7 |
| Episodes duration<br>(minutes) | 0.8 ± 0.1 | 0.71 ± 0.1 | 0.9 ± 0.1 |
| Frequency<br>(episodes/hour) | 33.1 ± 3.2 | 30.1 ± 1.6 | 32.1 ± 1.7 |
| **Light Sleep (LS)** |  |  |  |
| Number of episodes | 177.0 ± 16.9 | 185.7 ± 5.0 | 179.6 ± 16.6 |
| Episodes duration<br>(minutes) | 0.2 ± 0.0 | 0.2 ± 0.0 | 0.2 ± 0.0 |
| Frequency<br>(episodes/hour) | 47.7 ± 4.4 | 46.4 ± 1.2 | 44.9 ± 4.1 |
| **Slow Wave Sleep (SWS)** |  |  |  |
| Number of episodes | 133.9 ± 11.6 | 162.4 ± 7.9 | 148.4 ± 10.1 |
| Episodes duration<br>(minutes) | 0.5 ± 0.1 | 0.6 ± 0.0 | 0.6 ± 0.0 |
| Frequency<br>(episodes/hour) | 36.0 ± 2.9 | 40.6 ± 2.0 | 37.1 ± 2.5 |
| **Intermediate Stage (IS)** |  |  |  |
| Number of episodes | 20.7 ± 9.0 | 25.0 ± 8.0 | 14.8 ± 3.9 |
| Episodes duration<br>(minutes) | 0.2 ± 0.0 | 0.3 ± 0.0 | 0.3 ± 0.0 |
| Frequency<br>(episodes/hour) | 5.2 ± 2.3 | 6.3 ± 2.0 | 2.9 ± 0.9 |
| **REM Sleep** |  |  |  |
| Number of episodes | 18.3 ± 5.8 | 17.6 ± 2.3 | 10.4 ± 2.2 |
| Episodes duration<br>(minutes) | 1.0 ± 0.2 | 0.9 ± 0.1 | 0.9 ± 0.1 |
| Frequency<br>(episodes/hour) | 4.6 ± 1.3 | 4.4 ± 0.6 | 2.6 ± 0.6 |
| **Latency (minutes)** |  |  |  |
| NREM | 13.3 ± 3.7 | 6.2 ± 2.0 | 17.4 ± 5.5 |
| REM | 75.2 ± 9.6 | 75.1 ± 14.4 | 100.6 ± 22.1 |

Group differences were determined by repeated measures ANOVA.

**Figure 3.**
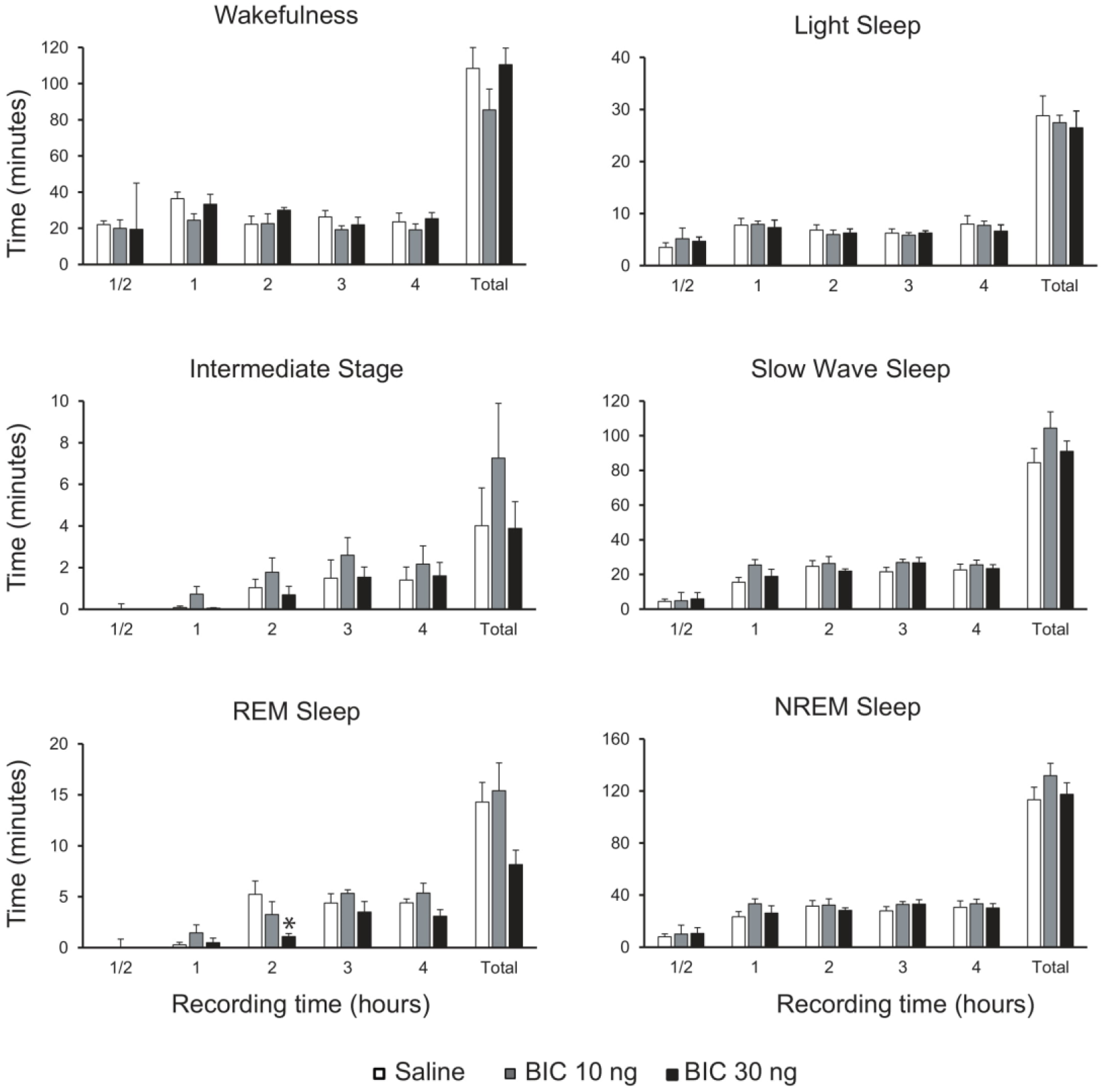
Effects of bicuculline microinjections into the mPOA on sleep and wake parameters. Graphic charts show the mean ± the standard error of the time spent in wakefulness, light sleep, slow wave sleep, no-REM sleep (NREM, light sleep + slow wave sleep), intermediate stage and REM sleep after local administration of saline and BIC (10 and 30 ng/ 0.2 µl) during the first half hour, each hour individually and the total recording time (four hours). Group differences were determined by one-way repeated measures ANOVA followed by Tukey as *post hoc*; Asterisk (*) indicates significant difference compared to control value.

The power spectrum profiles in different behavioral stages are shown in Figure 4 and Supplementary Figure S1, for the total recording time and the first half hour respectively, where it is evidenced that no differences were found in any frequency band power, cortex or behavioral stage among groups, neither in the total recording time nor in the first half hour.

**Figure 4.**
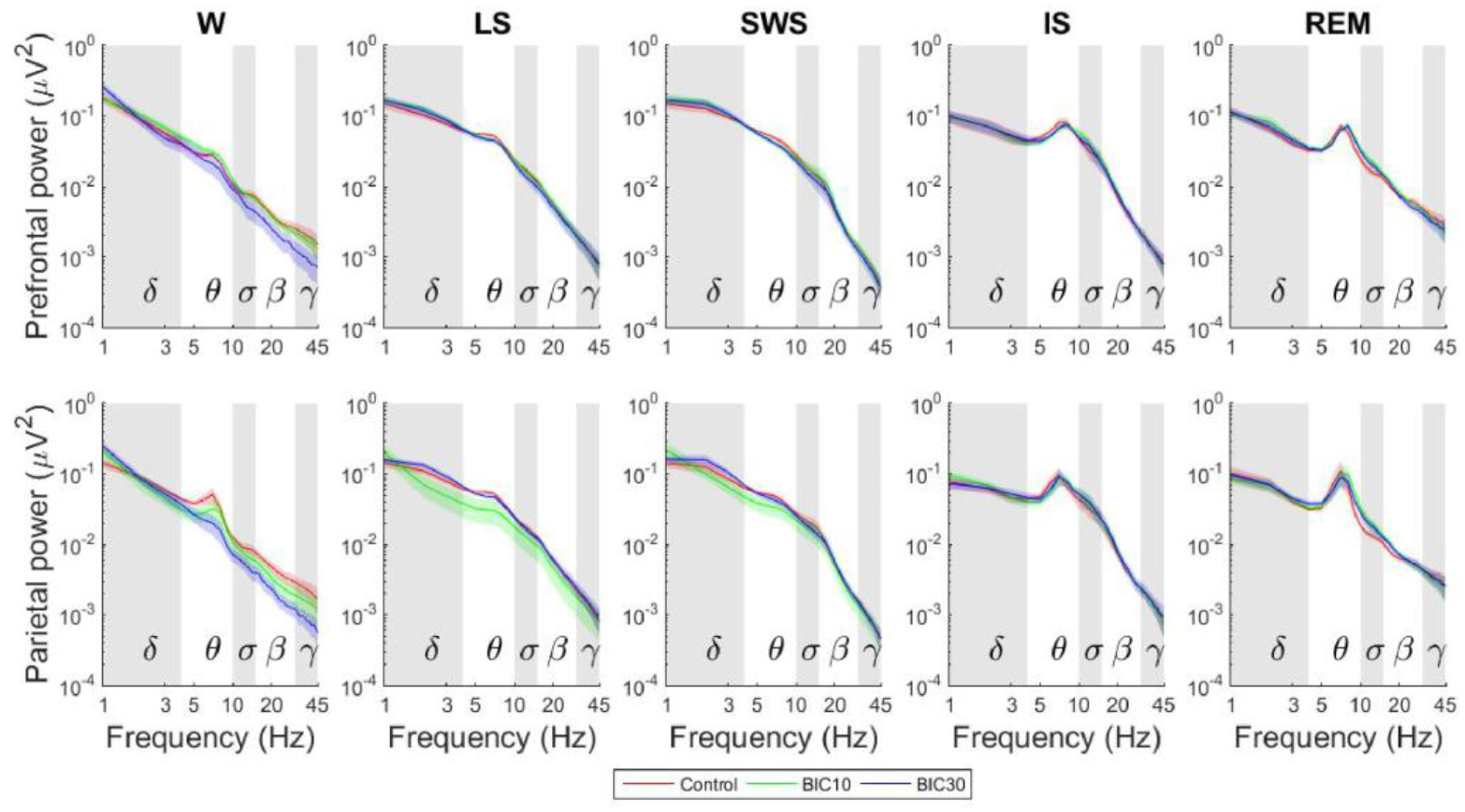
Mean relative power profile of each behavioral state during the entire recording time (4 hours) after bicuculline (BIC) treatment and vehicle within the mPOA. The graphs plot spectral power changes in the prefrontal and parietal regions for frequencies between 1 and 45 Hz during the different behavioral stages after BIC (10 and 30 ng/0.2 µl) or vehicle treatment. Thick and dark lines represent mean values and its correspondent standard error is represented in a shading of the same light color. Frequency bands are indicated by alternating horizontal-colored background of the graphs. Group differences were determined by one-way repeated measures ANOVA followed by Tukey as *post hoc*. No significant differences were found. W, wakefulness; LS, light sleep; SWS, slow wave sleep; REM, REM sleep.

### 3.2. Bicuculline effects on VLPO

#### 3.2.1. Maternal behavior

As shown in Figure 2, after BIC_30_ microinjection into the VLPO, the number of pups retrievings increased compared to control microinjections (T_8_ = 0.0, p = 0.011). Also, the number of mouthings augmented at this dose compared to saline group (T_8_ = 3.5, p = 0.042). However, the number of anogenital licking decreased at both doses compared to saline microinjections (BIC_10_: T_8_ = 0.0, p = 0.011; BIC_30_: T_8_ = 0.0, p = 0.017).

Most passive maternal behaviors were disrupted after the highest dose, and some of them with the lowest one (see Table 3). Specifically, the time hovering over the pups was reduced at both doses compared to vehicle values (BIC_10_: T_8_ = 3.0, p = 0.035; BIC_30_: T_8_ = 2.0, p = 0.025) as well as the total time in contact with the pups (BIC_10_: T_8_ = 0.0, p = 0.011; BIC_30_: T_8_ = 2.0, p = 0.025). The total time spent in nursing the pups over the 30-minutes test was slightly decreased after the highest dose compared to vehicle (T_8_ = 4.0, p = 0.049). Also, the latency to begin hovering over the pups (BIC_10_: T_8_ = 2.0, p = 0.025; BIC_30_: T_8_ = 3.0, p = 0.035) and nursing (BIC_10_: T_8_ = 3.0, p = 0.035; BIC_30_: T_8_ = 3.0, p = 0.035) were increased after BIC microinjection at both doses compared to vehicle.

**Table 3.** Maternal behavior after BIC and saline microinjections into the VLPO. Effects of bilateral microinjections of saline and BIC (10 and 30 ng/0.2 μl/side) into the ventrolateral preoptic area (VLPO) on maternal behavior.

|  | **VLPO** |  |  |
| --- | --- | --- | --- |
|  | **Control** | **BIC<sub>10</sub>** | **BIC<sub>30</sub>** |
| *Latency to (seconds):* |  |  |  |
| Reunion litter (sec) | 144.5 ± 233.0 | 189.0 ± 262.9 | 429.5 ± 345.9 |
| Hover over | 101.0 ± 58.9 | 508.5 ± 271.5* | 671.5 ± 120.6* |
| Nursing posture | 295.0 ± 82.0 | 604.0 ± 176.5* | 780.5 ± 176.8* |
| *Duration of (seconds):* |  |  |  |
| Hover over | 493.5 ± 113.1 | 186.0 ± 94.6* | 181.0 ± 77.0* |
| Nursing postures | 1077.5 ± 203.8 | 1075.0 ± 355.3 | 861.0 ± 583.0* |
| Contact with pups | 1668.0 ± 131.5 | 1267.5 ± 397.1* | 1111.5 ± 594.5* |
| Milk ejections (number) | 2.0 ± 0.25 | 1.0 ± 1.0 | 1.5 ± 1.1 |
| Litter weight gain | 4.8 ± 0.9 | 5.4 ± 0.8 | 3.7 ± 0.5 |

Group differences were determined by Friedman followed by Wilcoxon matched-pair signed-ranks test; significant differences compared to saline treatment are indicated by asterisks (\*, $p < 0.05$ ).

No other significant differences were found neither between each dose compared to vehicle group nor between BIC_10_ and BIC_30_.

Non-maternal behaviors (self-grooming and eating) were not affected by either doses of BIC in comparison to saline microinjections (see Table S1 in Supplementary data).

As a control procedure, we compared all the maternal behavior parameters between saline microinjection into the VLPO and mPOA; no significant differences were observed.

#### 3.2.2. Sleep

As depicted in Figure 5, statistical analysis over the total recording time showed significant differences in W time (F (2, 14) = 6.342, p=0.010), with an increase after BIC_30_ compared to BIC_10_ (p = 0.018) and control values (p = 0.023). Also, the total time spent in LS differ between groups (F (2, 14) = 6.302, p = 0.011), being reduced when comparing BIC_10_ (p = 0.034) and BIC_30_ (p = 0.014) with control group (see Figure 5). In addition, the duration of LS episodes varied among groups (F (2, 14) = 9.861, p = 0.002), where both BIC_10_ (p = 0.009) and BIC_30_ (p = 0.002 where significantly reduced compared to saline treatment (see Table 4).

**Table 4.** Sleep and waking effects after bicuculline (BIC) and saline microinjections into the ventrolateral preoptic area (VLPO). Effects of bilateral microinjections of saline and BIC (10 and 30 ng/0.2 μl/side) into VLPO on sleep parameters.

|  | **Control** | **BIC<sub>10</sub>** | **BIC<sub>30</sub>** |
| --- | --- | --- | --- |
| **Wakefulness** |  |  |  |
| Number of episodes | 143.9 ± 11.7 | 144.8 ± 9.8 | 148.1 ± 13.3 |
| Episodes duration<br>(minutes) | 0.66 ± 0.1 | 0.7 ± 0.1 | 0.8 ± 0.1 |
| Frequency<br>(episodes/hour) | 35.97 ± 2.9 | 36.2 ± 2.3 | 37.0 ± 3.1 |
| **Light Sleep (LS)** |  |  |  |
| Number of episodes | 203.3 ± 19.8 | 173.3 ± 11.6 | 169.0 ± 13.0 |
| Episodes duration<br>(minutes) | 0.2 ± 0.0 | 0.1 ± 0.0* | 0.1 ± 0.0* |
| Frequency<br>(episodes/hour) | 50.8 ± 4.9 | 43.3 ± 2.7 | 42.3 ± 3.0 |
| **Slow Wave Sleep (SWS)** |  |  |  |
| Number of episodes | 148.9 ± 11.7 | 160.5 ± 9.7 | 150.4 ± 8.3 |
| Episodes duration<br>(minutes) | 0.6 ± 0.1 | 0.7 ± 0.0 | 0.6 ± 0.0 |
| Frequency<br>(episodes/hour) | 37.2 ± 2.9 | 40.1 ± 2.3 | 37.6 ± 1.9 |
| **Intermediate Stage (IS)** |  |  |  |
| Number of episodes | 26.2 ± 4.5 | 28.1 ± 5.4 | 19.9 ± 4.8 |
| Episodes duration<br>(minutes) | 0.2 ± 0.0 | 0.2 ± 0.0 | 0.3 ± 0.0 |
| Frequency<br>(episodes/hour) | 6.6 ± 1.1 | 7.0 ± 1.3 | 5.0 ± 1.1 |
| **REM Sleep** |  |  |  |
| Number of episodes | 22.8 ± 2.6 | 24.9 ± 4.9 | 12.8 ± 2.8 # |
| Episodes duration<br>(minutes) | 0.7 ± 0.1 | 0.6 ± 0.1 | 0.9 ± 0.1 |
| Frequency<br>(episodes/hour) | 5.7 ± 0.5 | 6.2 ± 1.1 | 3.2 ± 0.6 # |
| **Latency (minutes)** |  |  |  |
| NREM | 9.3 ± 0.9 | 8.5 ± 1.5 | 18.1 ± 3.9 # |
| REM | 53.2 ± 8.8 | 60.7 ± 11.4 | 70.0 ± 13.9 |

Group differences were determined by repeated measures ANOVA followed by Tukey post hoc test; significant differences compared to saline treatment are indicated by asterisks (\*, $p < 0.05$ ); significant differences compared to BIC<sub>10</sub> treatment are indicated by numeral (#, $p < 0.05$ ).

**Figure 5.**
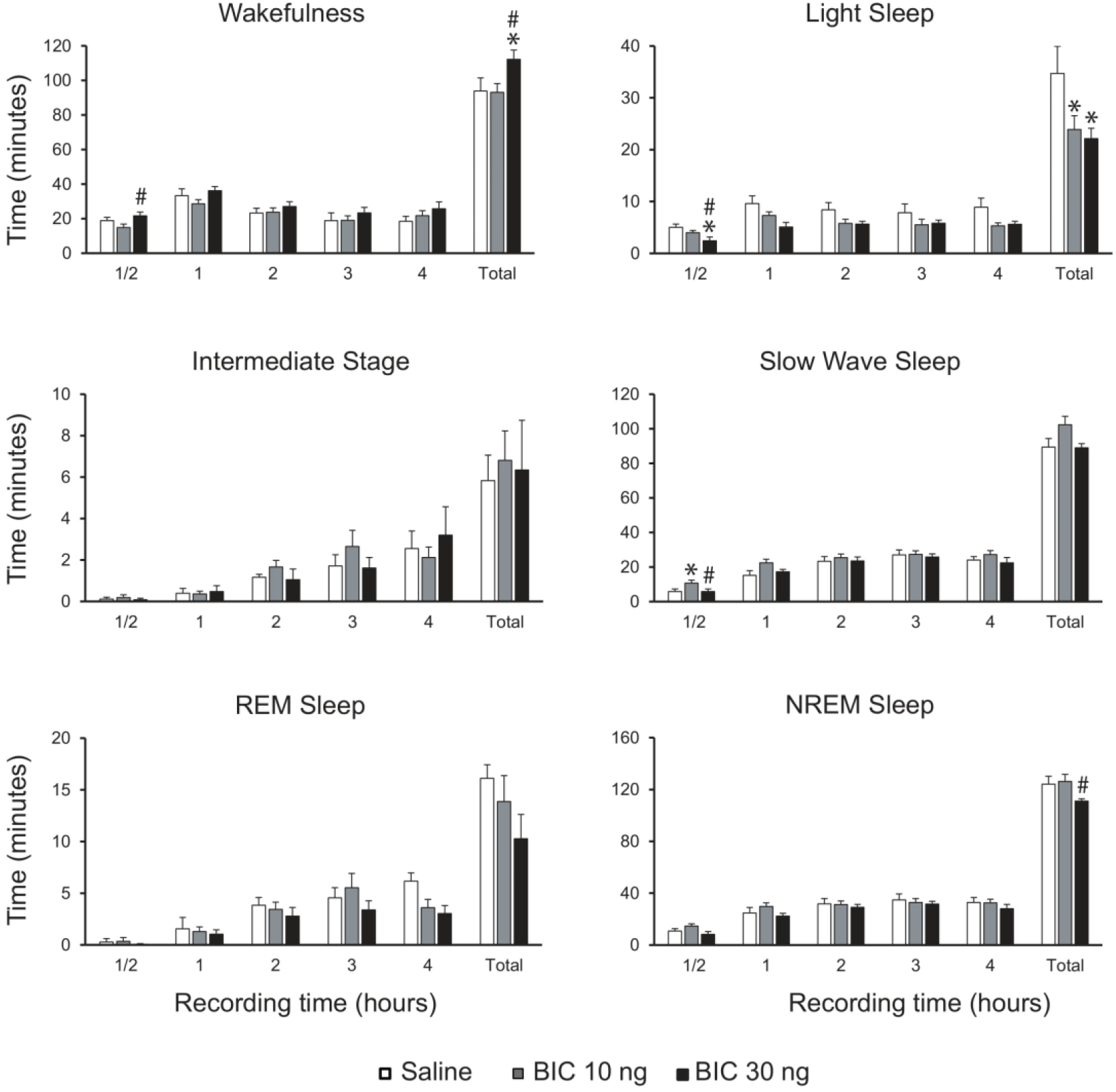
Effects of bicuculline microinjections into the VLPO on sleep and wake parameters. Graphic charts show the mean ± the standard error of the time spent in wakefulness, light sleep, slow wave sleep, no-REM sleep (NREM, light sleep + slow wave sleep), intermediate stage and REM sleep after local administration of saline and BIC (10 and 30 ng/ 0.2 µl) during the first half hour, each hour individually and the total recording time (four hours). Group differences were determined by one-way repeated measures ANOVA followed by Tukey as *post hoc*; Asterisk (*) indicates significant differences compared to control values; numeral (#) indicates significant differences compared to BIC10 ng.

The total time in SWS significantly differ among groups (F (2, 14) = 4.029, p=0.041). Although this parameter tended to increase in BIC_10_ group compared to BIC_30_ (p = 0.062) and saline treatment (p = 0.072), none of the *post hoc* analysis reached significant values.

The total time in NREM sleep differ between groups (F (2, 14) = 4.198, p = 0.037). Specifically, after BIC_30_ microinjection this time was reduced compared to BIC_10_ treatment (p = 0.045) and tend to be reduced when comparing to saline group (p = 0.087).

REM values over the total recording time tended to be different among groups but did not reach significant differences (F (2, 14) = 3.077, p=0.078). However, as shown in Table 4, the total number of REM episodes differ among groups (F (2, 14) = 4.380, p=0.033), being BIC_30_ reduced compared with BIC_10_ (p = 0.037) and had a slightly tendency to be reduced when compared to saline treatment (p = 0.091). In addition, the frequency of REM episodes per hour was also significantly different among groups (F (2, 14) = 4.380, p = 0.033), being significantly lower BIC_30_ compared to BIC_10_ treatment (p = 0.038).

During the first half hour, when maternal behavior was tested, the time spent in W differ among groups (F (2, 14) = 7.294, p = 0.006), where BIC_10_ treatment had less W compared to BIC_30_ (p = 0.005). In addition, the first half hour the time spent in LS differ among groups (F (2, 14) = 11.864, p = 0.000). Specifically, LS time was reduced after BIC_30_ compared to both saline (p = 0.000) and BIC_10_ treatment (p = 0.030). Furthermore, SWS also was affected by BIC treatment the first half hour (F (2, 14) = 5,128, p = 0.021), being increased after BIC_10_ compared to saline (p = 0.037) and BIC_30_ treatment (p = 0.037).

The mean power spectrum in different behavioral states during the total recording time and the first half hour were plotted in Figure 6 and Supplementary Figure S2 respectively. In particular, the theta frequency band in the prefrontal cortex during REM sleep in the total recording time was significant different among groups (F (2, 14 = 5.131, p = 0.021, see Figure 6). Specifically, BIC_30_ was significantly reduced compared to control (p= 0.049) and to BIC_10_ values (p = 0.009). No additional differences were found in any other frequency band power among groups at any behavioral state or cortex.

**Figure 6.**
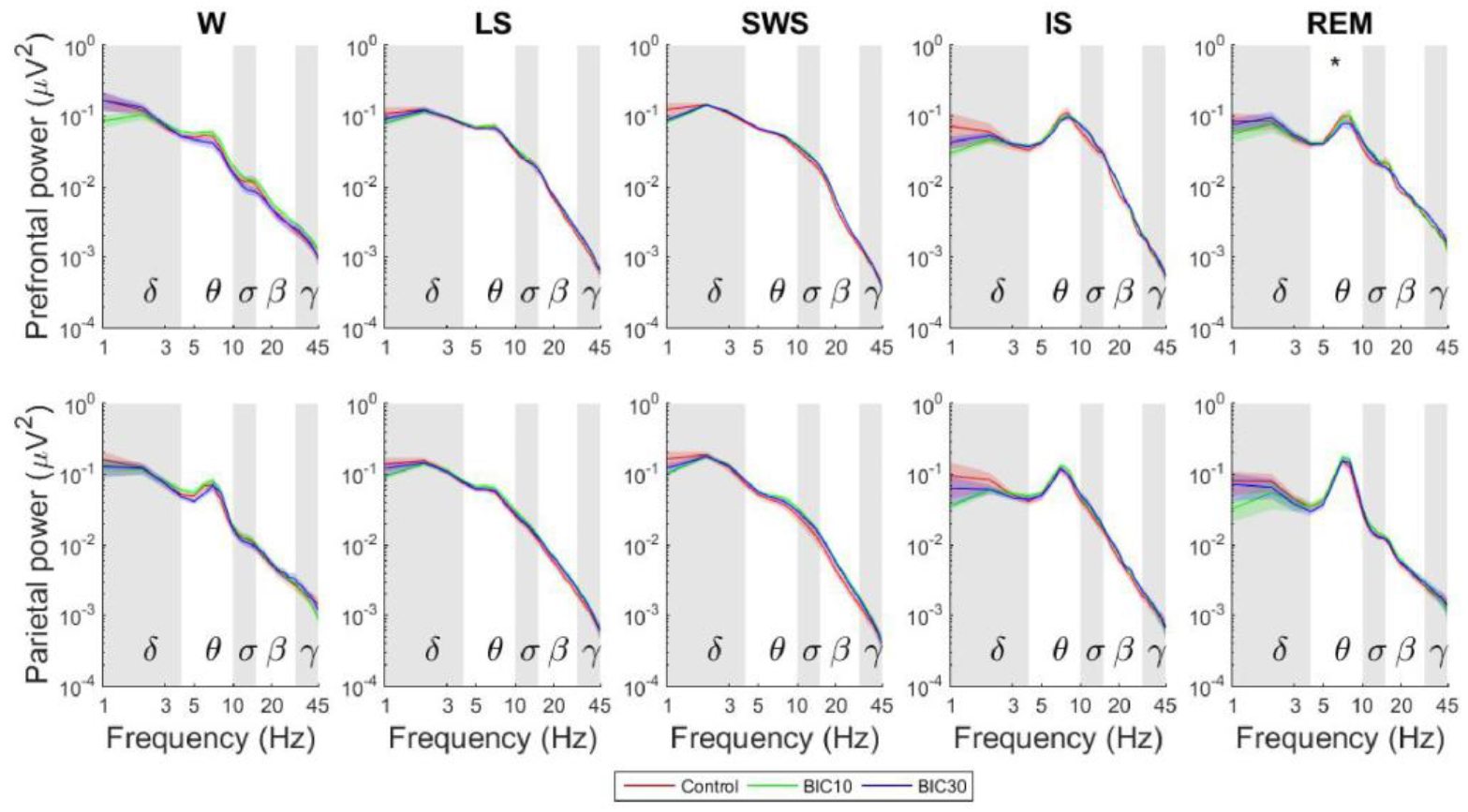
Mean relative power profile of each behavioral state during the entire recording time (4 hours) after bicuculline (BIC) treatment and vehicle within the VLPO. The graphs plot spectral power changes in the prefrontal and parietal regions for frequencies between 1 and 45 Hz during the different behavioral stages after BIC (10 and 30 ng/0.2 µl) or vehicle treatment. Thick and dark lines represent mean values and its correspondent standard error is represented in a shading of the same light color. Frequency bands are indicated by alternating horizontal-colored background of the graphs. Group differences were determined by one-way repeated measures ANOVA followed by Tukey as *post hoc*. Significant difference between BIC30 and control group is indicated by an asterisk (*, p < 0.05). W, wakefulness; LS, light sleep; SWS, slow wave sleep; REM, REM sleep.

As a control procedure, we compared all sleep parameters between saline microinjection into the VLPO and mPOA; no significant differences were observed in the total recording time or latencies in any of the behavioral stages.

## 4. Discussion

In the present study, we evaluated the regulation of sleep and maternal behavior after the microinjection of the GABA-A antagonist bicuculline into the mPOA and VLPO. Maternal behavior was similarly affected by BIC when applied either within the mPOA or VLPO. In addition, sleep and wakefulness values were modified when BIC was applied into the VLPO but almost unchanged by the same treatment within the mPOA.

### 4.1. mPOA and maternal behavior

The local infusion of BIC within the mPOA selectively enhanced some active components of maternal behavior, while reducing nursing behavior. Specifically, retrieving and nest building were enhanced after local delivery of BIC into the mPOA. In this sense, the GABA-A receptor agonist muscimol, microinjected in the mPOA produced a reduction of retrieving and nest building while leaving nursing intact [26]. A large body of evidence show that lesions or inactivation of mPOA [40, 41] and disruptions of its connection with the mesocorticolimbic system [42, 43] mainly affect the active components of maternal behavior such as the retrieving of the pups.

Although it was demonstrated the presence of GABA-A receptors within the mPOA [25], it is uncertain which are the neuronal phenotype and cellular localization within this area that we are antagonizing in the present report. It can be hypothesized that BIC may disinhibit either local inhibitory GABAergic interneurons or neurons that provide inhibitory input to other neural sites, allowing meaningful sensory stimulus coming from the pups activate the mPOA that project to different brain areas that stimulate maternal care of the pups [44]. In this sense, Lonstein et al. (2000) proposed that the neurons within mPOA that show elevated c-fos activity during the display of maternal behavior are either local inhibitory GABAergic interneurons or provide inhibitory input to other neural sites [45]. Otherwise, BIC can be acting directly into galaninergic neurons within the mPOA that have been demonstrated to be crucial for parenting behavior [46]. More studies are needed to assess if the impairing in retrieving is due to local mechanisms within the mPOA or to its ascending projections.

The reduction in the time devoted to nurse the pups after BIC injections may be related to the increased time invested in active maternal behavior. This result differs with that of Arrati et al (2006) showing that muscimol, microinjected in the mPOA did not affect nursing [26]. Although both drugs are GABA-A agents, and would expect opposite actions, differences in the dose or protocol may account for this discrepancy. In addition, maternal behavior is not a unitary process but rather a complex group of sequential behaviors that have their own sensory and neural determinants [40]. Specifically, it has been shown that nursing behavior is not primary controlled mPOA, but by brainstem-spinal areas [40, 41]. Therefore, it can be speculated that the reduction of the time devoted to nurse may be a consequence of the investment in the active care of the pups.

### 4.2. VLPO and maternal behavior

In an attempt to explore if there is an overlap in the neural networks related to sleep and maternal behavior, we evaluated maternal behavior after the blockade of GABAergic neurotransmission within the VLPO, a key area in sleep control. As expected, the increase in the time spent in wakefulness was accompanied by a decreased in nursing time without changes in milk ejections. The fact that active maternal behaviors that involve more motor activity (pup-retrieving and mouthing) increased while anogenital licking was reduced, favors the hypothesis that maternal modifications are a consequence of the difficulty to sleep. In addition, non-maternal behaviors such a self-grooming or eating were not modified. Taken together, these results suggest that the capability of mother rat to stay still in the nest is reduced, devoting her augmented waking time to the execution of maternal behaviors that do not require relative quiescence.

### 4.3. mPOA and sleep

Here we reveal almost no effect on sleep when BIC was microinjected into the mPOA nor in the EEG analysis in any of the behavioral stages. In agreement with our present data, Chari et al. (1995) found no impact on sleep after local delivery of GABA into the mPOA [47]. Nevertheless, the role of the mPOA in the regulation of sleep has been demonstrated using different techniques [2, 8, 48-54]. However, since the protocol and doses applied for the mPOA were the same as in the VLPO, the results suggest that the GABAergic processes that control sleep in the postpartum rat are weaker within the mPOA than in the VLPO. In this sense, it has been shown that there are sex-specific differences in GABAA receptors [55], and within the mPOA, the expression of these receptors has an important hormonal dependency [56-58]. In this sense, in regions of the forebrain and hypothalamus, the sensitivity of the GABA-A receptor to neurosteroid modulation has also been shown to vary significantly with hormonal state (for review, [59-61]). Besides, as the mPOA undergoes great anatomical changes during the postpartum period [11, 13, 62], its functional role in sleep could be modified. Thus, the hormonal fluctuations during lactation could modify the capability of BIC administration into mPOA to modify sleep.

### 4.4. VLPO and sleep

Since Sherin et al. (1996), the VLPO has been largely established as a main brain structure in the control of sleep process [5, 28, 63-65]. However, the current data shows that the decrease in GABAergic transmission within the VLPO promotes wakefulness and reduces sleep in postpartum females, without great modifications of the EEG activity. In this sense, recent evidence show that the lateral preoptic area triggers awakening from sleep and that the waking state is accompanied with an increased in the cortical activity [66]. In addition, recent reports evidenced that chemogenetic stimulation of glutamatergic neurons induce wakefulness [67, 68]. Thus, it can be postulated that BIC would directly disinhibit these glutamatergic neurons that promote wakefulness.

We also observed that perfusion of BIC_30_ into the VLPO decreased theta power in the prefrontal cortex during REM sleep. A possible explanation could be the regulation of the VLPO projection toward areas that desynchronize this rhythm, such as the medial raphe nucleus [69, 70].

### 4.5. Conclusions

In the present study, using BIC as a pharmacological tool to disinhibit the mPOA and VLPO of postpartum rats, we evaluate the role of the mPOA and VLPO on sleep and maternal behavior. The results show that BIC microinjections into the mPOA increased certain active maternal behaviors and decreased nursing without affecting sleep and wakefulness, probably as a result of changes in the mechanisms that control maternal behavior. In contrast, the same drug in the VLPO provoked an increase in some active maternal behaviors and a decrease in nursing postures together with an enhancement of wakefulness, suggesting that maternal behavior can be altered by modifying a sleep-promoting area.

## Acknowledgements

We thank MSc. Joaquin Gonzalez for his technical support in EEG analysis. This work was partially supported by “Programa de Desarrollo de Ciencias Básicas (PEDECIBA)”. All authors have seen and approved the manuscript, and it hasn’t been accepted or published elsewhere. The authors have no competing interests.

## Notes

### Competing Interest Statement

The authors have declared no competing interest.

