## Supplementary data for "Local administration of bicuculline into the ventrolateral and medial preoptic nuclei modifies sleep and maternal behavior in lactating rats"

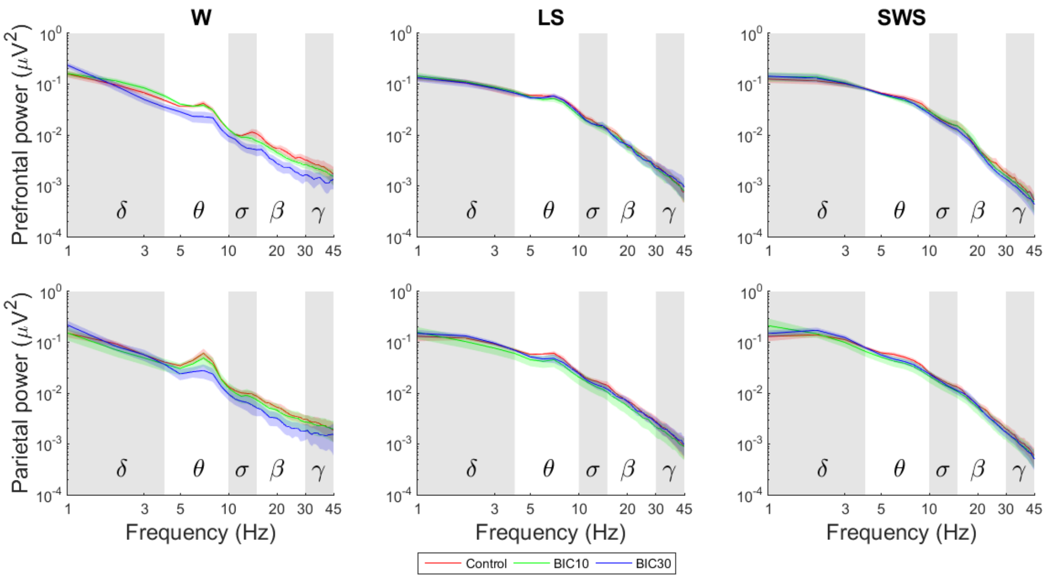


**Figure S1. Mean relative power profile of each behavioral state during the first half hour after bicuculline (BIC) treatment and vehicle within the mPOA.** The graphs plot spectral power changes in the prefrontal and parietal regions for frequencies between 1 and 45 Hz during the different behavioral stages after BIC (10 and 30 ng/0.2 µl) or vehicle treatment. Thick and dark lines represent mean values and its correspondent standard error is represented in a shading of the same light color. Frequency bands are indicated by alternating horizontal-colored background of the graphs. Group differences were determined by one-way repeated measures ANOVA followed by Tukey as *post hoc.* W, wakefulness; LS, light sleep; SWS, slow wave sleep; REM, REM sleep.

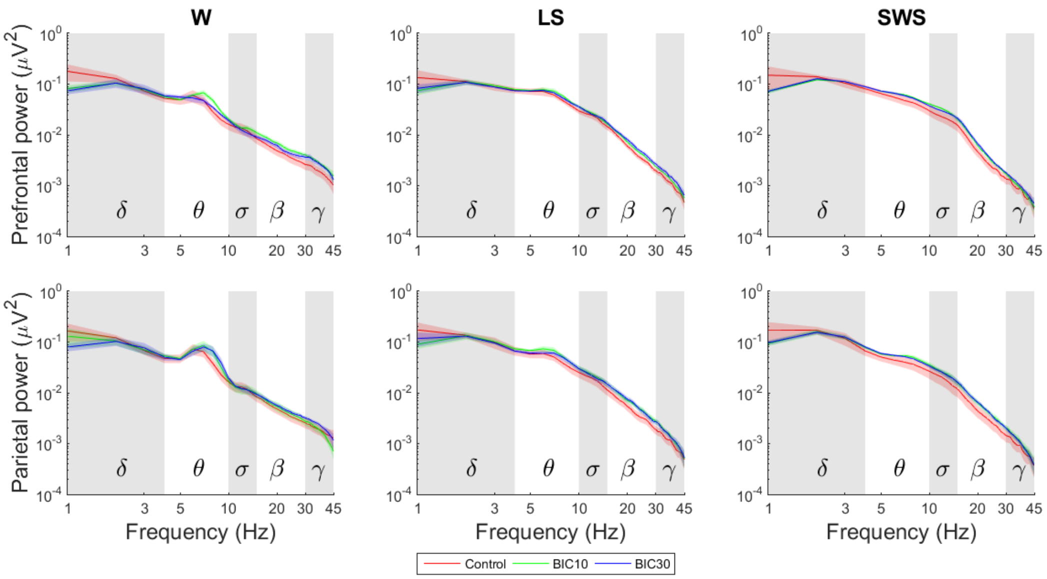


**Figure S2. Mean relative power profile of each behavioral state during the first half hour after bicuculline (BIC) treatment and vehicle within the VLPO.** The graphs plot spectral power changes in the prefrontal and parietal regions for frequencies between 1 and 45 Hz during the different behavioral stages after BIC (10 and 30 ng/0.2 µl) or vehicle treatment. Thick and dark lines represent mean values and its correspondent standard error is represented in a shading of the same light color. Frequency bands are indicated by alternating horizontal-colored background of the graphs. Group differences were determined by one-way repeated measures ANOVA followed by Tukey as *post hoc.* W, wakefulness; LS, light sleep; SWS, slow wave sleep; REM, REM sleep.

**Table S1. Non-maternal behavior after BIC and saline treatment into the mPOA and VLPO.** Effects of bilateral microinjections of saline and BIC (10 and 30 ng/0.2 μl/side) into the medial preoptic area (mPOA) and the ventrolateral preoptic area (VLPO) on eating and self-grooming during the 30-minute maternal behavior test.

| | **Control** | **$BIC_{10}$** | **$BIC_{30}$** |
|---|---|---|---|
| ***mPOA*** | | | |
| Self-grooming | 3.0 ± 1.3 | 6.0 ± 4.0 | 5.0 ± 1.5 |
| Eating | 0.0 ± 0.0 | 0.0 ± 0.0 | 0.0 ± 1.0 |
| ***VLPO*** | | | |
| Self-grooming | 5.0 ± 1.5 | 4.5 ± 2.4 | 4.0 ± 4.0 |
| Eating | 0.0 ± 0.0 | 0.0 ± 0.5 | 4.5 ± 9.0 |

Group differences were determined by Friedman test; no significant differences were found among groups.
